# Wild-type and mutated ß-catenin differently repress *RND3/RHOE* expression in hepatocellular carcinoma

**DOI:** 10.1101/2025.10.16.682743

**Authors:** Sara Basbous, Sandra Sena, Camille Dantzer, Véronique Neaud, Léo Piquet, Florence Grise, Frédéric Martins, Christine Varon, Sabine Gerbal-Chaloin, Alexandre Favereaux, Sabine Colnot, Valérie Lagrée, Clotilde Billottet, Violaine Moreau

## Abstract

**Background & Aims:** Tumor development and progression are mainly driven by oncogenic mutations but are also regulated by physical factors, such as applied forces or microenvironment stiffness. Through its structural and transcriptional functions, ßcatenin is a key factor that acts on both aspects to promote liver tumorigenesis, leading to hepatocellular carcinoma (HCC) development. However, the mechanisms by which these two functions regulate downstream targets remain poorly understood. Herein, we describe Rnd3, also called RhoE, an atypical member of the Rho GTPase family, as a common target of both functions of ß-catenin. We previously demonstrated that *RND3* expression is downregulated in HCC, which correlates with intrahepatic metastasis. Yet, a molecular understanding of how Rnd3 expression is dysregulated in cancer is largely missing.

**Approach & Results:** Using human HCC samples and cultured cell lines, we demonstrate that Rnd3 expression is regulated by ß-catenin pathways, regardless of their mutational status. Both the transcriptional and the structural activity of ß-catenin repress the expression of *RND3*. Indeed, we found that wild-type ß-catenin suppresses *RND3* transcription through the Hippo pathway, whereas oncogenic ß-catenin downregulates *RND3* expression through miRNA targeting its 3’UTR.

**Conclusion:** Rnd3 may constitute a key protein involved in the transcriptional program driven by oncogenic ß-catenin in HCC and as a mediator of the mechanosensitive response associated with cell-cell adhesion.

## INTRODUCTION

Tumor development and progression are mainly driven by oncogenic mutations but are also strongly influenced by physical factors, such as applied forces, cellular geometry or microenvironment stiffness. Indeed, intra- and intercellular communications coordinate various cellular activities including proliferation and migration through signal transduction pathways. ß-catenin is a key effector of these inside-out and outside-in mechanotransduction events (Astudillo, 2020). It plays a dual role in cells: i) as a mediator of adhesion and ii) as a transcriptional co-regulator of Wnt signaling. In normal epithelial cells, ß-catenin mainly localizes at the plasma membrane where it binds the cytoplasmic domain of cadherin adhesion receptors to maintain mechanosensitive adherens junctions. The extracellular adhesive activity of cadherins is linked to the underlying actin cytoskeleton *via* the actin-binding protein, α-catenin. In the absence of Wnt signaling, excess of cytoplasmic ß-catenin is phosphorylated by CK1 and GSK3ß and degraded by the ubiquitin-proteasome system. In contrast, upon Wnt signal, these kinases are inhibited and stabilized ß-catenin translocates to the nucleus where it binds DNA-binding factors, such as lymphoid enhancer factor-T-cell factor (LEF-TCF), to coordinate the activation or repression of target genes. Thus, in normal cells, ß-catenin is endowed with both structural and transcriptional roles. Through these functions, ß-catenin is involved in many key processes. As an essential intermediate of the Wnt signaling pathway, ß-catenin plays a key role in determining cell fate during embryonic development. In adult organs, Wnt signaling continues to play indispensable roles in tissue homeostasis, cell renewal, and regeneration (Valenta et al., 2012).

On the dark side, inappropriate activation of the Wnt/ß-catenin pathway has been largely implicated in carcinogenesis. In hepatocellular carcinoma (HCC), which is the main primary malignancy of the liver, aberrant transcriptional activation of the Wnt-targets occurs mainly through constitutively active ß-catenin mutations present in 18-41% of tumors (Cavard et al., 2008). Transcriptional programs and signatures linked to the oncogenic ß-catenin in HCC have been widely established and numerous ß-catenin target genes were described (Dantzer et al., 2024a). Herein, we identified the RhoGTPase Rnd3/RhoE as a novel target of ß-catenin signaling pathways in liver cancer.

Rnd3 (also called RhoE) belongs to the Rnd sub-family in the RhoGTPase family. Members of this sub-family (Rnd1-3) are considered as atypical GTPases because they lack any detectable GTPase activity and remain in the constitutively active GTP-bound form. Consequently, as Rnd proteins are not regulated by the typical GTP/GDP cycle, they are tightly controlled at transcriptional and post-transcriptional levels. Despite this atypical regulation, Rnd3 has been implicated in functions commonly regulated by Rho GTPases such as remodeling of the actin cytoskeleton and in many basic cellular processes such as cell proliferation, differentiation, survival, motility and adhesion (Paysan et al., 2016)(Riou et al., 2010). An increasing number of studies describe the alteration of Rnd3 expression in tumors and suggest that Rnd3 is a key gene in tumor progression (Basbous et al., 2020). Related to HCC, others and we demonstrated that *RND3* expression is down-regulated in tumoral tissues when compared to normal liver and that its low expression significantly correlates with intrahepatic metastasis and poor prognosis (Ma et al., 2013)(Grise et al., 2012)(Luo et al., 2012). We further demonstrated that Rnd3 down-regulation increases HCC cell invasion by promoting epithelial-mesenchymal transition (Grise et al., 2012). We recently described that the loss of Rnd3 induces reversible senescence (Basbous et al., 2022) and entosis, a mechanism of cell death *via* cell cannibalism (Basbous et al., 2024), in tumor hepatocytes. However, a molecular understanding of how Rnd3 expression is dysregulated in cancer is largely missing.

In this study, we demonstrate that Rnd3 expression is regulated by ß-catenin pathways. Rnd3 expression is significantly repressed in ß-catenin-mutated HCC compared to non-mutated HCC and negatively correlates with classical positive targets of ß-catenin in the liver. We found that ß-catenin silencing restores the expression of Rnd3 in HCC cell lines. Using the HepG2 model with dual ß-catenin knockdown (Gest et al., 2023), we found that *RND3* expression is indirectly repressed by both the transcriptional and the structural activity of ß-catenin. Indeed, wild-type (WT) ß-catenin inhibits *RND3* transcription through the Hippo pathway. In contrast, oncogenic ß-catenin represses *RND3* expression through miRNA targeting its 3’UTR. Thus, Rnd3 may represent a key protein involved in both WT and oncogenic ß-catenin signaling programs.

## MATERIALS AND METHODS

### Cell lines and primary hepatocyte culture

Human Hepatoblastoma (HB) cell line, HepG2, and human HCC cell lines, Hep3B, Huh7, SNU398 and SNU475 were purchased from American Type Culture Collection. The H2P HCC cell line and its metastatic counterpart H2M were a kind gift from Pr. Guan (Hu et al., 2004). HB cell line Huh6 was generously provided by Dr. Perret (Cochin Institute, Paris). All cell lines were maintained at 37°C in a humidified 5% CO_2_ atmosphere in the appropriate medium supplemented with 10% fetal calf serum (FCS), 100 U/mL penicillin, and 100 μg/mL streptomycin. HepG2, Huh7 and Hep3B cell lines were grown in Dulbecco’s modified Eagle’s medium (DMEM: 4.5 g/l glucose, glutamax, pyruvate). SNU398 and SNU475 cells were cultured in RPMI 1640 (4.5 g/l glucose) supplemented with L-glutamine. Huh6 cells were cultured in DMEM (1 g/l glucose) with L-glutamine and pyruvate; H2P and H2M cells in F12 medium. Cell line authentication was performed using short tandem repeat analysis, and absence of mycoplasma contamination in cell culture media was tested every week. The HepG2 shßcat MUT cell line expressing a shRNA specifically targeting mutated ß-catenin in a doxycycline-inducible manner, was generated previously (Dantzer et al., 2024b). These cells were cultured in the same conditions as parental HepG2 cells. shRNAs were induced with 1 μg/mL doxycycline. Fresh primary human hepatocytes were obtained from Biopredic SA (Rennes, France) and cultured following the provider’s instructions.

### Cell treatments

The GSK3 inhibitor CHIR99021 and the proteasome inhibitor MG132 (Sigma Aldrich) were used at 3 μM for 24h and 10 μM for 6h respectively. Human Wnt3a and R-spondin3 (RSPO3) were used at a final concentration of 100 ng/ml for 24h.

### Transfections

Gene knockdown was performed using small interfering RNAs (siRNAs) and lipofectamine RNAi max. Cells were seeded on day 1 in 6-well plates at a density of 300.000 cells in 2 mL of complete DMEM per well. Two rounds of transfection were performed: the first on day 1 (on non-adherent cells) and the second on day 3 (on adherent cells). For each transfection, a mixture of 1.5 μL of siRNA (from a 20 μM stock), 5 μL of Lipofectamine RNAi max, and 500 μL of Opti-MEM was prepared and added to the cells. If multiple siRNAs were transfected simultaneously, the amount of Lipofectamine was proportionally increased. The control siRNA corresponds to All Stars negative control from Qiagen. Cells were harvested on day 5 for RNA or protein extraction. For miRNA overexpression, miRNA (2μL from a 20μM stock) was co-transfected with siRNA during the second round of the transfection (day 3) and by adding an additional 5 μL of lipofectamine. The references or sequences of the siRNAs and miRNAs used are listed in supplemental Table 1.

To study the impact of *RND3* 3’untranslated region (3’UTR), human *RND3* 3’UTR (+1001 to +2807) was cloned in pLenti-UTR-GFP-Blank using EcoRI and XhoI sites to generate pLenti-RND3 3’UTR-GFP. pLenti-UTR-GFP-Blank (m014) was from Applied Biological Materials (abm). Lentiviral particules were produced by the Vect’UB platform (TBM core facility, Bordeaux). HepG2 cells were seeded in 6-well plates, then transduced after 24h at a multiplicity of infection of 5, either with a lentiviral plasmid containing the 3’untranslated (3’NT) part of Rnd3 coupled to GFP, or with a control plasmid encoding GFP without the 3’NT sequence. Transduced cells were selected using puromycin treatment (1 μg/mL).

For overexpression of YAP, plasmids were obtained from Addgene: pcDNA-Flag-Yap1 (addgene#18881; (Levy et al., 2008)) and pCMV-flag-S127A-YAP (addgene#27370; (Zhao et al., 2007)) encodes respectively, WT or constitutively activated human Yap1. DNA transfections were performed in HepG2 cells using Lipofectamine 3000 transfection reagent (ThermoFischer) according to manufacturer’s protocol. Control cells were empty vector transfected cells.

### Luciferase reporter assay

On day 4 after transfection of cells with siRNA targeting the different forms of ß-catenin, cells were detached and seeded on 96-well plates. Co-transfection was then performed with 64 ng of Rnd3 Firefly luciferase reporter plasmid (pGL4.10-Rnd3 promoter), previously published in (Piquet et al., 2018) and 25 ng of a constitutively active thymidine kinase (TK) promoter Renilla luciferase reporter plasmid. After 24h of incubation at 37°C, cells were lysed and analyzed using Dual-Luciferase® Reporter Assay (Promega). Firefly luciferase activity was normalized to Renilla luciferase activity to consider the variations in transfection efficiency. Thus, the ratio Firefly/Renilla was calculated and used for analysis.

### Real-time PCR

Quantitative RT-PCR (qRT-PCR) of 120 HCCs (57 from the “microarray” series and 63 new samples) and 28 nontumor liver samples was performed in duplicate, using TaqMan^®^ Low Density Array and the ABI PRISM^®^ 7900HT System (Applied Biosystems) as described previously (Boyault et al., 2007). For other analyses, RNA was collected from cultured cells using the Trizol reagent (Invitrogen) or RNA Isolation kit (Macherey-Nagel), according to manufacturer’s protocol. cDNA was synthesized from 2 μg of total RNA with Maxima® First Strand cDNA Synthesis Kit (Fermentas) using a mix of oligo(dT)_18_ and random hexamer primers. Aliquots of cDNA (2 to 30 ng according to target mRNA) were then subjected to PCR amplification on a StepOnePlus Real-Time PCR system (Applied Biosystems) with specific forward and reverse oligonucleotide primers. The SYBR^®^ Green SuperMix for iQ™ (Quanta Biosciences, Inc.) was used with the following PCR amplification cycles: initial denaturation, 95°C for 10 minutes, followed by 40 cycles with denaturation, 95°C for 15 seconds and annealing-extension, 60°C for 1 minute. A dissociation curve was generated to verify that a single product was amplified. Gene expression results were first normalized to internal control r18S. Relative levels of expression were calculated using the comparative (2-^ΔΔCT^) method. All primers used for qRT-PCR are listed in supplemental Table 2.

### Western-blot analysis

Total proteins were extracted from cells using RIPA lysis buffer (0.1% SDS, 1% NP40, 0.15 M NaCl, 1% sodium deoxycholate, 25 mM Tris HCl pH 7.4) supplemented with protease and phosphatase inhibitors (Roche). Proteins were then denaturated in Laemmli buffer (Bio-Rad) at 95°C for 5 min. 40 μg of protein extract was loaded on 10% polyacrylamide gels (TGX Stain-Free FastCast, Bio-Rad). Membranes were blocked with 5% BSA-TBST (5% BSA Sigma, TBS 1x Euromedex, 0,1% Tween20 Sigma) for 30 min and incubated with primary antibodies (diluted in 5% BSA-TBST) overnight at 4°C. The following primary antibodies were used: ß-catenin (1:2000, BD Bioscience, clone 14, #610154), Rnd3 (1:1000, clone 4, Cell signaling, #3664), GFP (1:1000, Abcam, #ab290), ß-actin (1:2000, Sigma Aldrich, #A2066), GAPDH (1:3000, Santa Cruz, #sc-25778). Membranes were then incubated with secondary antibodies (1:5000 diluted in 5% BSA-TBST) for 30 min. The following secondary antibodies were used: IRDye 680CW conjugated goat anti-rabbit IgG (H&L) (LI-COR), IRDye 800CW conjugated goat anti-mouse IgG (H&L) (LI-COR). Acquisitions were performed using the ChemiDoc Imaging System (Bio-Rad). Intensities were measured using the ImageLab software (Bio-Rad).

### Patient samples

Human HCC samples were described previously (Grise et al., 2012). As previously described, all samples came from resected or explanted livers with HCC of patients treated in Bordeaux from 1992-2005. Fragments of fresh tumor and non-tumor liver tissues (taken at a distance of at least 2 cm from the tumor) were immediately snap-frozen in liquid nitrogen and stored at -80°C. RNA or proteins were extracted as described (Boyault et al., 2007). HCCs used as the Affymetrix hybridization set (57 HCCs) and the qRT-PCR validation set (63 HCCs) were described (Boyault et al., 2007). The characteristics of HCCs used for the immunoblot analysis (27 HCCs) were indicated in Grise *et al*. (2012).

### miRNome analysis

Total RNA extraction was performed using the Direct-zol RNA Microprep kit (Zymo Research, R2062). RNA quantity was measured after extraction using a DS-11 spectrophotometer (Denovix) and RNA integrity was checked with the 2100 Bioanalyzer (Agilent). SmallRNA-seq libraries were then prepared using the QIAseq miRNA Library kit (QIAGEN, 331502) and QIAseq miRNA 96 index Kit IL UDI-F (QIAGEN, 331665) according to manufacturer’s protocol. Briefly, undiluted 3’ adapters were first ligated on 100ng of total RNA. Then, undiluted 5’ adapters were ligated and RNA retro-transcribed with undiluted RT primers to generate double strand cDNA. A final PCR of 16 cycles was performed to amplify libraries and barcode samples with unique dual indices. An additional purification step allowed to obtain 200-300 bp fragments. Profiling libraries was performed with the LabChip GX Touch HT Nucleic Acid Analyzer (Revvity), with HT DNA NGS 3K Reagent Kit (Revvity, CLS960013) using the HT DNA X-Mark Chip (Revvity, CLS144006). Then, libraries were quantified by qPCR using the LightCycler 480 II (Roche) with the KAPA Library Quantification Kit (Kapa, KK4854). Finally, libraries were sequenced at the PGTB facility (https://doi.org/10.15454/1.5572396583599417E12) with the NextSeq 1000/2000 P2 Reagents (100 Cycles) v3 (Illumina, 20046811) using the NextSeq 2000 sequencer platform (Illumina). Single read sequencing (72 bp) resulted in an average of 22 million single reads by sample. The small RNAseq data have been deposited to the EBI repository with the dataset identifier: E-MTAB-15638.

### Statistical tests

Data are presented as the mean ± SD of at least three independent experiments. Statistical tests were performed using GraphPad Prism software version 10. Normality was assessed using Shapiro-wilk test and comparisons between groups were made using Student’s t-test or the Mann Whitney U test. For the correlation, Pearson’s t test was used. *P* values are indicated as such: * *P* < 0.05; ** *P* < 0.01; *** *P* < 0.001; **** *P* <0.0001; *ns*, not significant.

## RESULTS

### Rnd3 expression is down-regulated in human ß-catenin-mutated HCCs

Others and we have previously demonstrated that Rnd3 expression is down-regulated in HCC at both protein and mRNA levels (Ma et al., 2013)(Grise et al., 2012)(Luo et al., 2012). Data extracted from the GEPIA database (Tang et al., 2019) combining expression in 529 samples confirmed that *RND3* expression is significantly repressed in HCC (Fig. 1A). As genetic alterations in the *RND3* gene are rare events in HCC (Basbous et al., 2020), we searched for transcriptional mechanisms that may repress *RND3* expression. We first used the Boyault et al transcriptional data that includes 57 HCC samples and 5 pooled non-tumor tissues, and analyzed the level of *RND3* in the six groups of HCC (G1-G6) according to J. Zucman-Rossi’s classification (Boyault et al., 2007). These groups overlap proliferative (G1 to G3) and non-proliferative (G4 to G6) tumors as described by others (Dantzer et al., 2024a). We found that *RND3* is significantly down-regulated in all HCC groups but highly significantly in subgroups G5 and G6 (Fig. 1B), enriched in *CTNNB1*-mutated HCCs. We indeed found that *RND3* down-expression correlated significantly with *CTNNB1* mutations in this cohort (Fig. 1C). On the other hand, we found that *RND3* expression level was not correlated to *TP53* mutations in HCC (Fig. S1A). We further analyzed the correlation of *RND3* expression with known positive ß-catenin transcriptional targets, as *LGR5* and *GLUL*, encoding respectively G-protein coupled receptor 49 (Gpr49) and Glutamine Synthetase (GS). Consistently, we found that *RND3* expression correlates negatively with the expression of these classical ß-catenin targets in HCC (Fig. 1D). Quantitative RT-PCR performed on the same sample set and on a second independent set of 63 tumors, demonstrated that *RND3* mRNA expression was significantly lower in ß-catenin-mutated HCCs when compared to ß-catenin-non-mutated tumors (Fig. 1E). This tendency extended to the protein level, where Rnd3 protein was found significantly downregulated in HCC expressing high level of GS (HCC GS(+)) (Fig. S1B). As *CTNNB1* is also mutated at a high level, i.e. up to 80%, in hepatoblastoma, we further checked for *RND3* expression in these pediatric liver tumors. Data extracted from Cairo et al. transcriptomic analysis (Cairo et al., 2008) show that *RND3* mRNA level is strongly decreased in tumors compared to non-tumoral liver, regardless of C1 or C2 subgroups classification (Fig. S1 C-D). Moreover, this downregulation was even more pronounced in ß-catenin-mutated tumors (Fig. S1E). Thus, these data suggest that *RND3* expression is repressed by activated or mutated ß-catenin in liver tumors.

**Figure 1:**
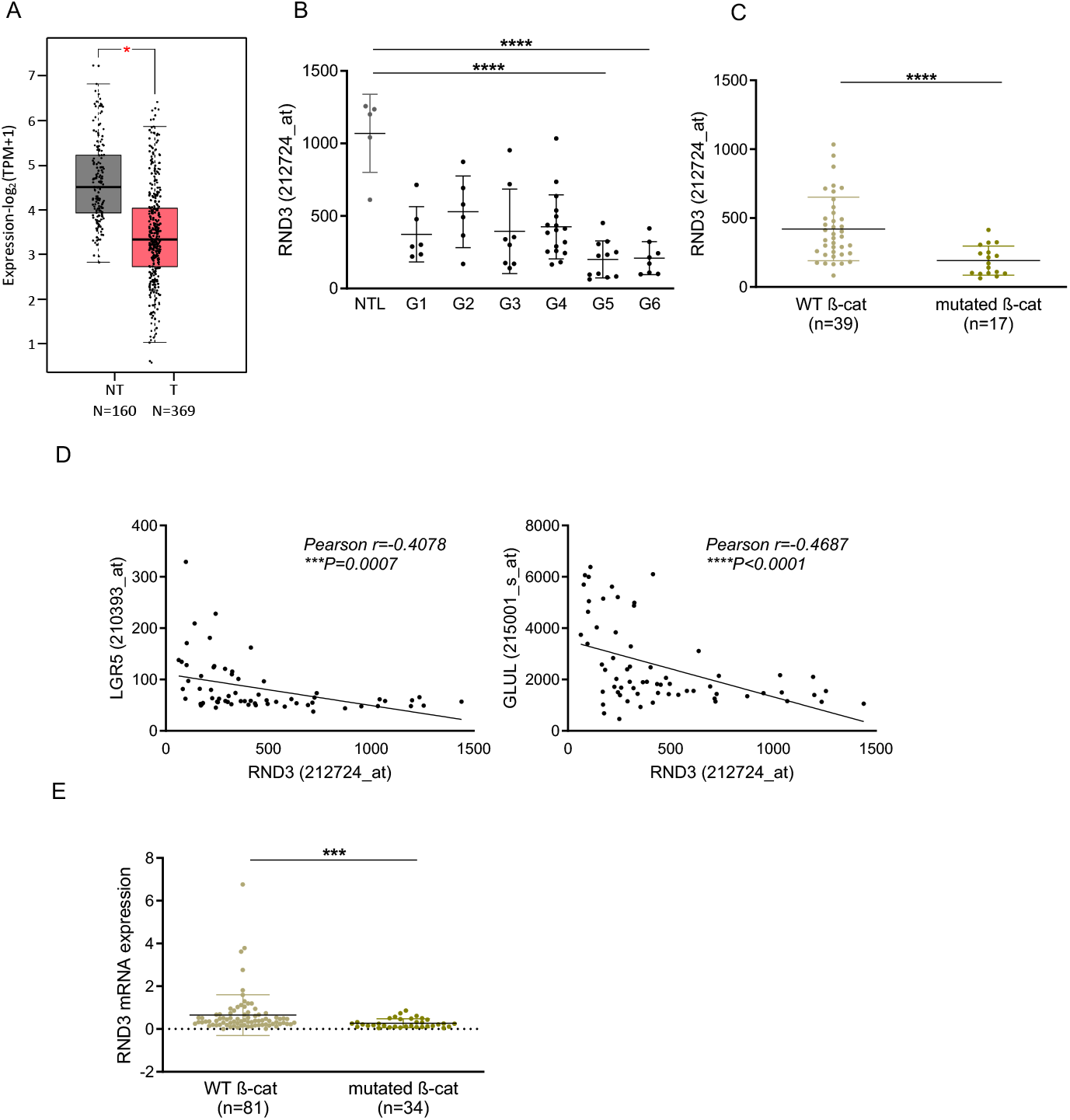
Rnd3 expression is downregulated in human ß-catenin-mutated HCCs. **(A)** *RND3* gene expression in non-tumor (NT, n=160; gray histogram) and tumor (T, n=369; red histogram) human liver samples using RNA-seq data from GEPIA (TCGA and GTEx). **(B)** *RND3* gene expression from transcriptomic data of the Boyault cohort, including 5 non-tumor (NTL) tissues and 57 HCC samples classified into six subgroups (G1-G6) according to J. Zucman-Rossi’s classification (Boyault et al., 2007). **(C)** *RND3* gene expression in human HCC with wild-type (WT, n=39) versus mutated β-catenin (n=17) from the Boyault et al. cohort. **(D)** Correlation of expression between the *RND3* gene and ß-catenin positive target genes, *LGR5* and *GLUL* (n=65 patients). **(E)** qRT-PCR validation of *RND3* mRNA expression on WT (n=81) and ß-catenin-mutated (n=34) tumors. Results are expressed as Mean ±SD. Statistical test was determined using Mann Whitney, Student’s t-test or Pearson correlation test. \**P*<0.05; \*\*\*\**P*<0.0001.

### Rnd3 expression is altered by ß-catenin signaling in liver tumor cells

As we published earlier, Rnd3 protein expression is altered in liver tumor cell lines compared with primary hepatocytes (see Fig. 2A in Grise *et al*., 2012). Notably, we observed that the lowest Rnd3 protein expression was found in ß-catenin-mutated cell lines specifically in HepG2 and Huh6 cell lines. We further confirmed that *RND3* mRNA levels were markedly reduced in both HepG2 and Huh6 cells (Fig. 2A) where, as expected, both ß-catenin transcriptional targets, *LGR5* and *AXIN2* were highly expressed (Fig.2A).

**Figure 2:**
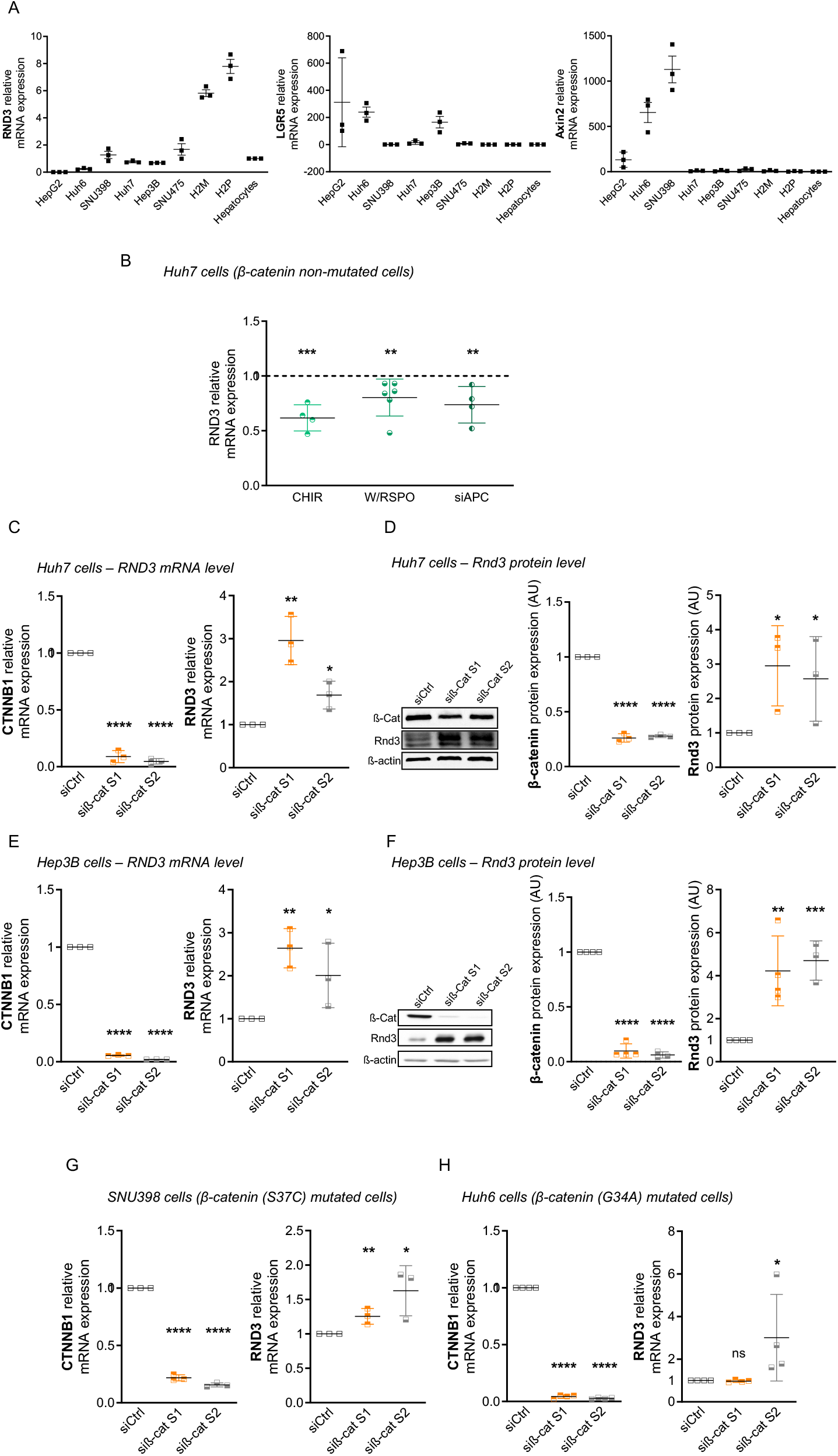
Rnd3 expression is regulated by ß-catenin signaling in liver tumor cells. **(A**) mRNA expression of *RND3*, and ß-catenin target genes *LGR5* and *AXIN2* in different liver tumor cell lines. Cell lines with ß-catenin mutated (HepG2, Huh6, SNU398) or wild-type ß-catenin (Huh7, Hep3B, SNU475, H2M, H2P) were analyzed and the mRNA expression was normalized to normal hepatocytes (n=3). **(B)** *RND3* mRNA expression in Huh7 cells after activation of ß-catenin pathway through inhibition of GSK3β using CHIR99031 (CHIR) or treatment with wnt3a/spondin3 (W/RSPO) or APC knockdown with siRNA (siAPC). The results are shown relative to their respective control treated cells, respectively DMSO, PBS or siCtrl. **(C-F)** Huh7 (C, D) and Hep3B (E, F) cells were transfected with two different siRNA targeting ß-catenin (Si β-cat S1 and Si β-cat S2). *CTNNB1* (ß-catenin) and *RND3* expressions were assessed at both mRNA expression (C, E) and protein levels (D, F). **(G-H)** In cells with point mutations in ß-catenin, SNU398 (G) and Huh6 (H), *RND3* expression was evaluated by qRT-PCR after ß-catenin knockdown. Results are representative of at least three independent experiments and expressed as Mean ±SD. Statistical significance was determined using Mann Whitney, Student’s t-test. *P<0.05; **P<0.01; ***P<0.001; \*\*\*\**P*<0.0001.

In order to study whether the activation of the Wnt/ß-catenin pathway can alter *RND3* mRNA expression in HCC cells, we applied various treatments to *CTNNB1-*non-mutated Huh7 cells: inhibition of GSK3ß with the CHIR99031 inhibitor, application of Wnt3a/spondin3 and APC inhibition using siRNA. All three treatments led to a decrease of *RND3* mRNA expression in Huh7 cells (Fig. 2B) demonstrating that the activation of Wnt/ß-catenin pathway represses the expression of *RND3*. We next performed the reverse experiment by depleting ß-catenin using two independent siRNAs targeting ß-catenin in *CTNNB1*-non mutated Huh7 and Hep3B cell lines. These siRNA showed an efficient knockdown of ß-catenin expression and its targets *AXIN2* and *LGR5* (Fig.2C-F, and Fig.S2A-B). Interestingly, ß-catenin depletion led to a significant increase of Rnd3 expression at the mRNA and protein levels (Fig.2C-F) suggesting that WT ß-catenin negatively regulates Rnd3. Furthermore, using the same siRNAs in *CTNNB1*-mutated SNU398 and Huh6 cell lines, we demonstrated that the inhibition of mutated ß-catenin significantly increases the expression of Rnd3 (Fig.2G-H and Fig. S2C-D). Thus, ß-catenin represses *RND3* expression regardless of its mutational status in HCC cells. Altogether, these results highlight an unexplored way of *RND3* regulation and establish *RND3* as a novel negative target of ß-catenin.

### *RND3* expression is regulated differently by structural and transcriptional functions of ß-catenin

As previously described, ß-catenin has two functions, as a transcriptional co-regulator of Wnt signaling and as a mediator of cell-cell adhesion. The observation that *RND3* mRNA levels are up-regulated upon ß-catenin silencing in cells carrying ß-catenin mutations or not, led us to hypothesize that both the transcriptional and structural functions of ß-catenin may repress *RND3* expression. To test this hypothesis, we made use of the so-called “dual ß-catenin knocked-down HepG2 model”, that we recently characterized and published (Gest et al., 2023). This model allows to dissociate the structural and the transcriptional activities of β-catenin, carried respectively by the WT and mutated *CTNNB1* alleles. We thus analyzed *RND3* expression using three different siRNA depleting WT, mutated or both forms of ß-catenin in HepG2 cells. As expected, the expressions of classic transcriptional ß-catenin targets such as *AXIN2* and *LGR5*, were strongly repressed upon silencing of mutated or both ß-catenin (Fig. S3). In contrast, *RND3* repression was released upon removal of either the WT or the mutated ß-catenin (Fig. 3A). Indeed, Rnd3 expression is upregulated by 10-fold when either form was silenced and about 25-fold when both simultaneously depleted (Fig. 3A), suggesting that WT and mutated ß-catenin repress *RND3* expression *via* two different pathways. These new findings were confirmed at the protein level (Fig. 3B). Since Rnd3 protein levels are very low in HepG2 cells, we additionally treated cells with the proteasome inhibitor MG132 to enhance Rnd3 protein detection. The Rnd3 protein was significantly upregulated following knock-down of WT ß-catenin and slightly upon mutated ß-catenin. Consistent with results obtained at mRNA levels, the silencing of both forms led to an additive effect (Fig. 3B), meaning that Rnd3 expression is independently repressed by WT and mutated ß-catenin in HepG2 cells. Altogether, our data demonstrated that across all liver tumor cell types tested, Rnd3 expression is repressed by ß-catenin pathways. Moreover, we found that ß-catenin can mediate this repression through both its transcriptional and structural functions.

**Figure 3:**
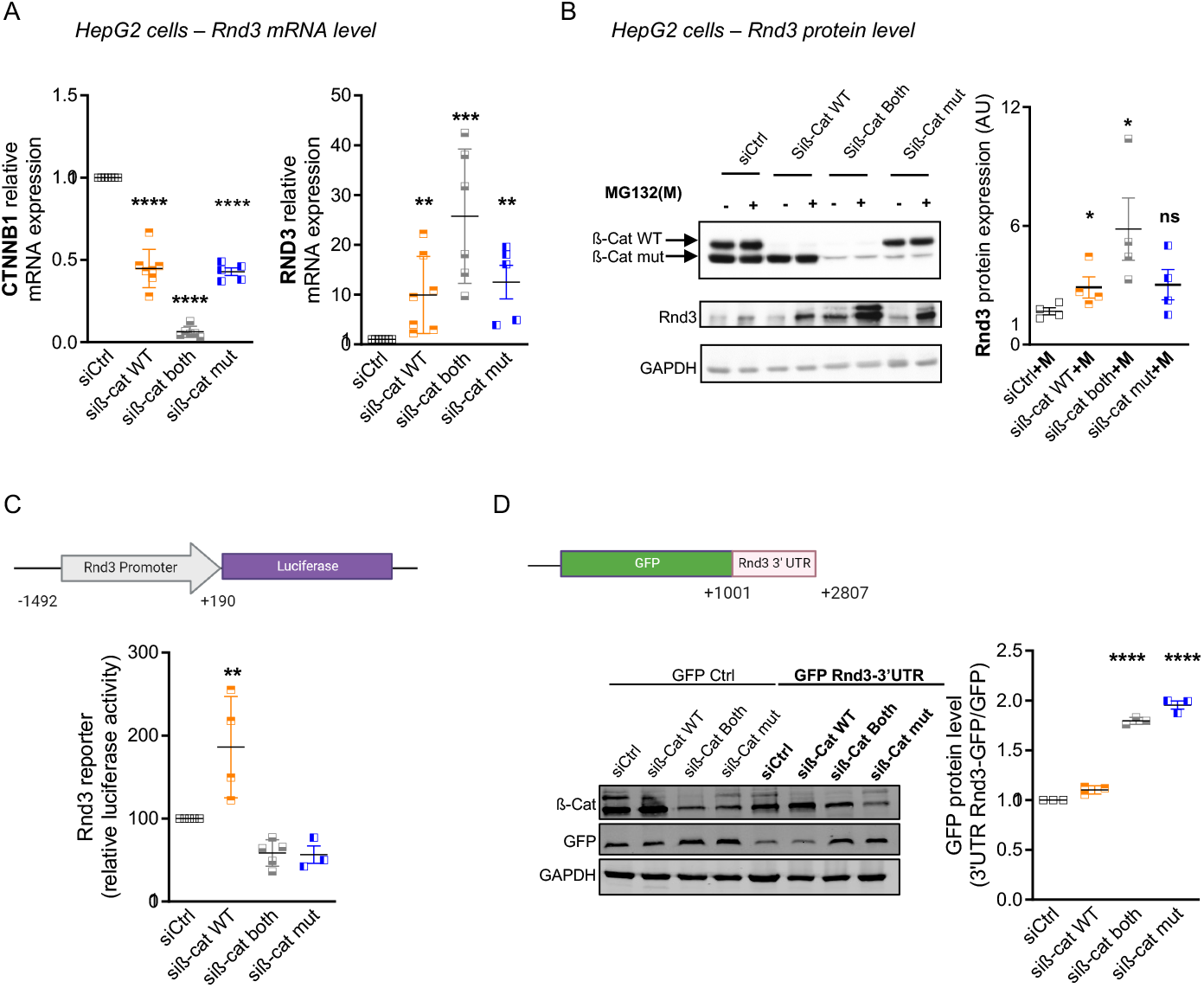
Rnd3 expression is differently regulated by the wild-type and mutated ß-catenin. **(A)** *CTNNB1* and *RND3* mRNA expression were analyzed by qRT-PCR in HepG2 cells upon inhibition of wild-type (siß-cat WT), mutated (siß-cat mut) or both (siß-cat both) ß-catenin. **(B)** Rnd3 expression was evaluated at the protein level by western-blot after inhibition of WT, mutated or both β-catenin with siRNAs and treatment with or without the proteasome inhibitor MG132 (M). The graph shows quantification of Rnd3 protein normalized to GAPDH. **(C)** Luciferase reporter assay was performed in HepG2 cells co-transfected with Rnd3 promoter-luciferase and siRNAs targeting different forms of ß-catenin. **(D)** *RND3* mRNA stability was analyzed using a construct expressing GFP under the control of *RND3* 3’UTR. GFP expression was evaluated by western-blot after ß-catenin inhibition with the different siRNAs. Results are representative of at least three independent experiments and expressed as Mean ±SD, Student’s t-test. *P<0.05; **P<0.01; ***P<0.001; \*\*\*\**P*<0.0001.

To further study how the ß-catenin regulates *RND3* expression, we performed gene reporter assays with either the *RND3* promoter upstream luciferase or the *RND3* 3’untranslated region (3’UTR) coupled to GFP. The impact of ß-catenin extinction on the activity of the Rnd3 promoter or 3’UTR was evaluated after plasmid transfection and treatment with the three different siRNAs targeting ß-catenin in HepG2 cells. Unexpectedly, depletion of WT ß-catenin, but not the mutated form, led to an increase in the activity of *RND3* promoter (Fig. 3C) suggesting that the WT ß-catenin represses *RND3* expression transcriptionally through its promoter. Conversely, when assessing the impact of ß-catenin silencing on *RND3* 3’UTR, only the inhibition of mutated ß-catenin increases the expression of GFP meaning that the mutated form regulates *RND3* expression through its 3’UTR (Fig. 3D). Altogether, these results demonstrated two different ways of *RND3* regulation, i.e. a transcriptional regulation by the WT ß-catenin and a post-transcriptional regulation by the mutated ß-catenin.

### The structural activity of ß-catenin represses *RND3* expression via YAP/TEAD pathway

The analysis of *RND3* promoter revealed several consensus binding sites for various transcription factors such as LEF/TCF, but also TEAD transcription factors. To study how WT ß-catenin represses *RND3* expression, we again used the “dual ß-catenin knocked-down HepG2 model”. Previous transcriptomic analysis of this model highlighted an upregulation of genes associated with the mechanosensitive Hippo pathway upon depletion of WT ß-catenin (see Fig. S2E-F in Gest *et al*., 2023). We confirmed these results by quantitative PCR and demonstrated a significant increase of several target genes of the pathway such as *ANKRD1, CYR61, AXL, AMOTL2, and ARHGAP29* (Fig. 4A). The expression of other members of the pathway, including the TEAD transcription factors (*TEAD1* and *TEAD2*) and *MST1/2*, were also significantly upregulated upon WT ß-catenin silencing in HepG2 cells (Fig. 4B). Based on these findings, we hypothesized that WT ß-catenin decreases the expression of Rnd3 by negatively regulating *TEAD1* and/or *TEAD2*. To test this hypothesis, we transfected HepG2 cells with siRNA targeting the WT form of ß-catenin either alone or in combination with siRNA depleting *TEAD1* or *TEAD2* or both (Fig. 4C-E). As showed before, WT ß-catenin silencing led to increased *RND3* expression. Interestingly, this expression decreases significantly when we co-depleted the WT ß-catenin with either TEAD1 or TEAD2 (Fig. 4C-D, Fig. S4A-B). We observed the same results when ß-catenin, TEAD1 and TEAD2 are simultaneously inhibited (Fig. 4E, Fig. S4C). As YAP and TAZ are key effectors of the Hippo pathway, we asked whether their inhibition, in combination with that of WT ß-catenin could counteract the effect observed with WT ß-catenin alone, similar to what was seen upon TEAD1/2 inhibition. Indeed, the co-inhibition of WT ß-catenin and YAP using siRNA reversed the effect of the WT ß-catenin alone (Fig. 4F, Fig. S4D). We have not observed any effect using the siRNA targeting TAZ suggesting that WT ß-catenin acts *via* YAP, but not TAZ, to regulate the expression of *RND3* (Fig. S4E). Moreover, we found that the overexpression of YAP increases significantly *RND3* expression, as it does for its transcriptional target *ANKRD* (Fig. 4G). In addition, the activation of YAP using its constitutively active mutated form, S127A, increases much more the *RND3* mRNA expression compared to the overexpression of WT YAP (Fig. 4G). According to these results, analysis of HCC samples without ß-catenin mutations from the TCGA cohort revealed a correlation between *RND3* expression and the YAP/TAZ pathway target genes such as *CYR61, AXL, AMOTL2* and *ARHGAP29* (Fig. 4H). Thus, our findings demonstrate that WT ß-catenin represses *RND3* expression by modulating the Hippo pathway.

**Figure 4:**
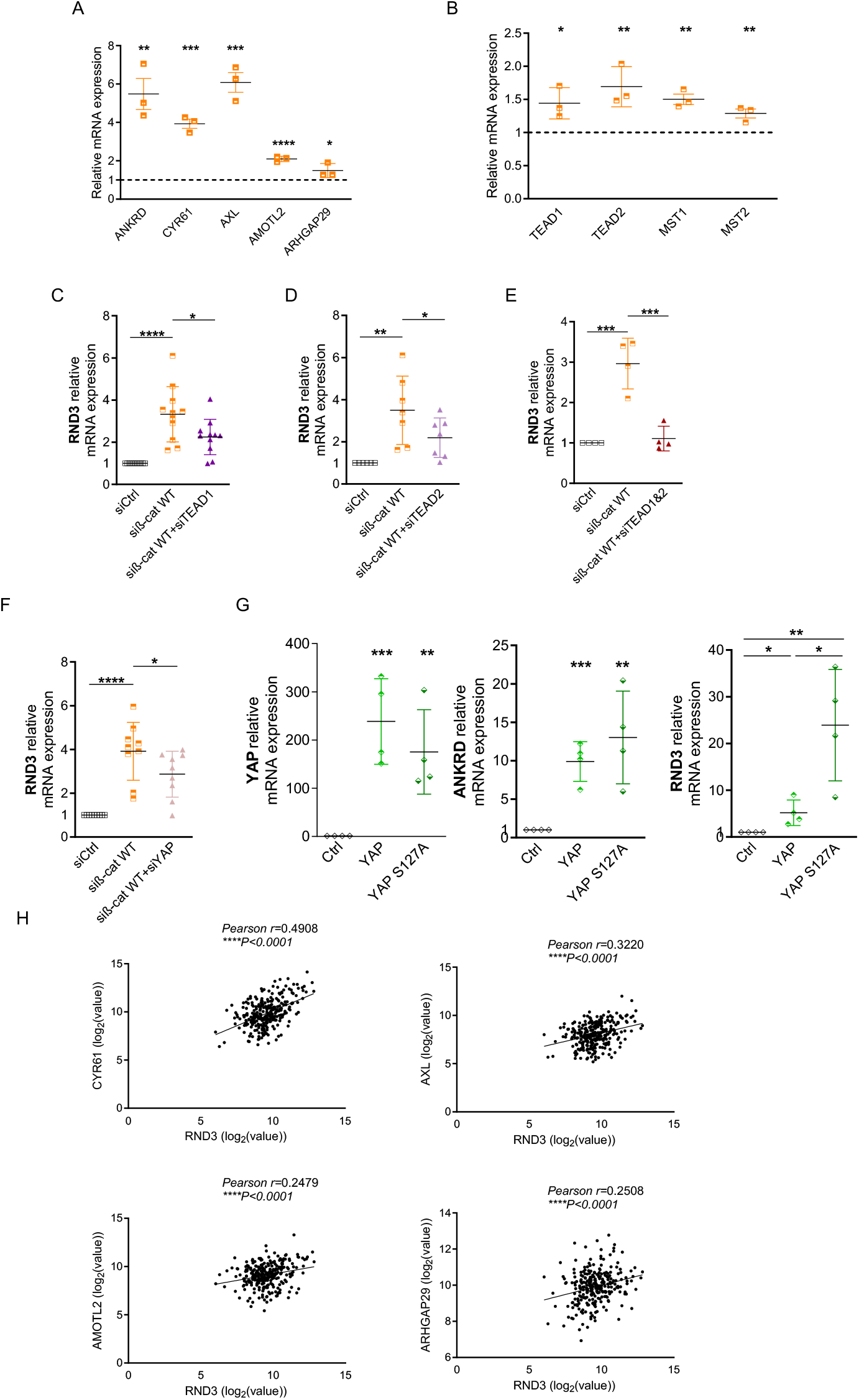
The structural activity of ß-catenin regulates *RND3* via YAP/TEAD pathway. **(A-B)** HepG2 cells were transfected with siRNA targeting the WT ß-catenin, and the mRNA expression of TEAD target genes *ANKRD, CYR61, AXL, AMOTL2, ARHGAP29* (A) and genes in the Hippo pathway (transcription factors (*TEAD1, TEAD2*) and *MST1, and MST2* (B) were evaluated by qRT-PCR. **(C-F)** HepG2 cells were transfected with siRNA against WT form of ß-catenin alone or in combination with siRNA targeting *TEAD1* (C) or *TEAD2* (D) or both (E) or YAP (F) and the mRNA expression of *RND3* was evaluated by qRT-PCR. Results are representative of at least three independent experiments. **(G)** HepG2 cells were transfected with WT or mutant (S127A) YAP and the expression of YAP and its targets *ANKRD* and *RND3* were evaluated. **(H)** Correlation between *RND3* and the YAP/TEAD target genes *CYR61, AXL, AMOTL2*, and *ARHGAP29* in human HCC patients with non-mutated ß-catenin using TCGA dataset (n=277 patients). Results are presented as Mean ±SD, Student’s t-test or Pearson correlation test. *P<0.05; **P<0.01; ***P<0.001; ****P<0.0001.

### The transcriptional activity of ß-catenin represses *RND3* expression via miR-512-3p

We then sought to understand how the oncogenic form of ß-catenin regulates *RND3* expression via its 3’UTR. In order to identify miRNAs whose expression is altered upon mutated ß-catenin silencing, we performed the miRNAome of our dual ß-catenin knocked-down HepG2 model, that we improved with doxycycline-inducible shRNAs (Dantzer et al., 2024b). Analysis of miRNA specifically altered upon mutated ß-catenin silencing compared to the control condition revealed 175 differentially expressed miRNAs, of which 85 were upregulated and 90 were downregulated (Fig. 5A, supplemental Table 3). Among those significantly downregulated miRNAs, 7 were predicted by miRDB to target *RND3* 3’UTR (Fig. 5A-B). The most significantly down-regulated, miR-512-3p, has been previously implicated in the regulation of *RND3* expression in prostate cancer cells (Rao et al., 2018). Interestingly, we found that the overexpression of miR-512-3p reverses the effect on *RND3* mRNA due to ß-catenin depletion in HepG2 cells (Fig. 5C). These results showed that the mutated ß-catenin represses *RND3* expression *via* its effect on miR-512-3p. To further validate this finding, gene expression data from the TCGA cohort were used and showed significant negative correlations between *RND3* and miR-512-3p expressions in patients with HCC harboring ß-catenin mutations (Fig. 5D).

**Figure 5:**
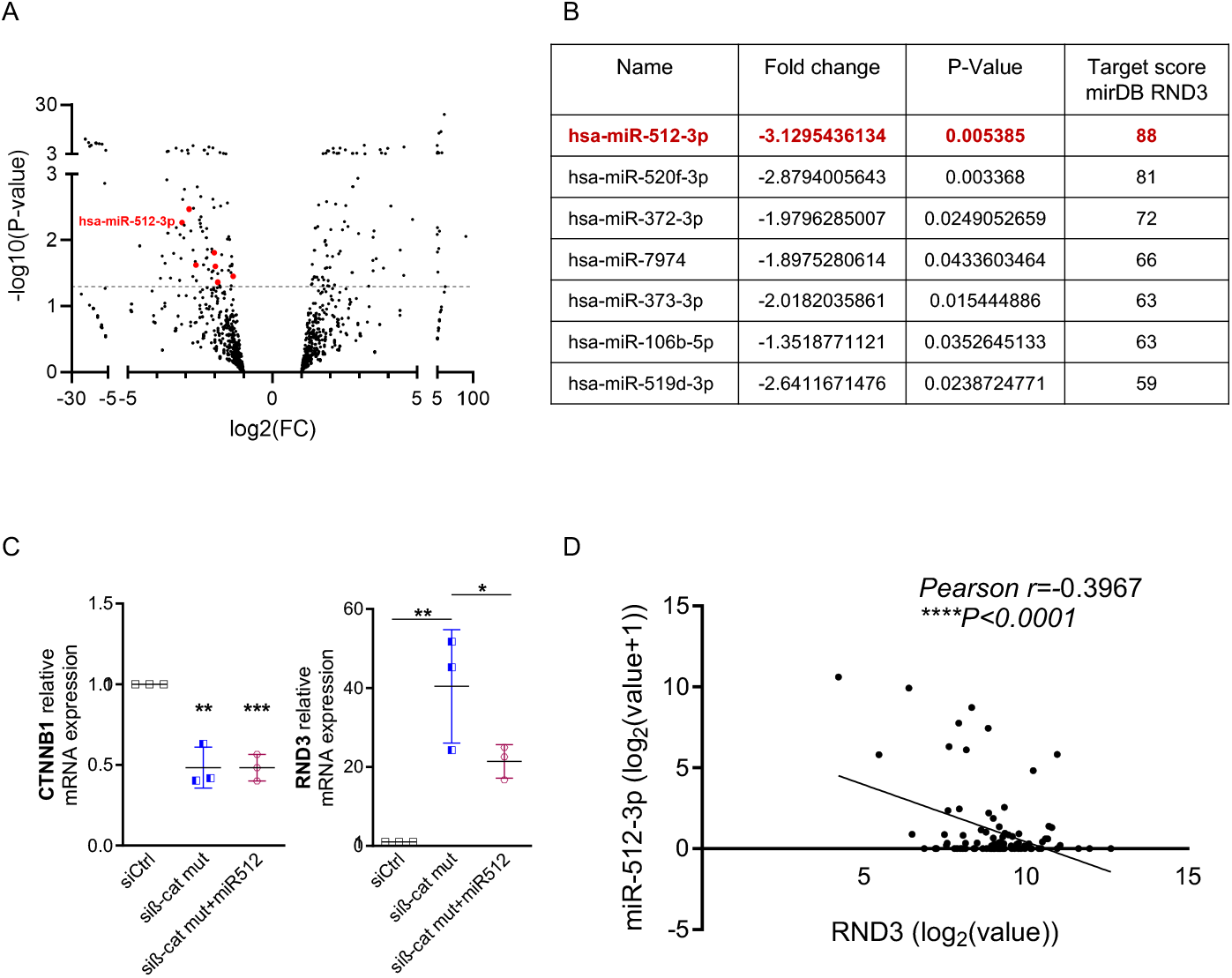
The mutated form of ß-catenin represses *RND3* expression *via* miR-512-3p. **(A)** miRNAome analysis of HepG2 expressing doxycycline-inducible shRNA against the mutated form of ß-catenin or a control shRNA, showing differentially expressed miRNAs. miRNAs predicted to target *RND3* 3’UTR are highlighted in red. hsa-miR-512-3p is one among the downregulated miRNAs after downregulation of mutated ß-catenin (n=3). **(B)** Table listing significantly downregulated miRNAs and predicted to target *RND3* 3’UTR, based on miRDB (https://mirdb.org/). **(C)** Expression of *RND3* mRNA expression in HepG2 cells after inhibition of mutated ß-catenin with or without miR-512-3p overexpression (n=3). **(D)** Correlation between miR-512-3p and *RND3* expression in human HCC patients with mutated ß-catenin using TCGA dataset (n=93 patients). Results are expressed as Mean ±SD, Student’s t-test. *P<0.05; \*\**P*<0.01; \*\*\**P*<0.001.

## DISCUSSION

ß-catenin is a key protein involved in liver development and physiology, where it maintains liver zonation and regulates liver function. However, it is also a true driver of liver carcinogenesis, found mutated in up to one third of human HCCs and more than 80% of hepatoblastomas. With ß-catenin described as an undruggable target, it remains important to better understand the ß-catenin pathway and identify downstream targets that may play key role in development or progression of liver tumors. Here, we have identified *RND3* as one such target. In most tumors, ß-catenin mutations are present in a heterozygous state, meaning that mutated and WT alleles co-exist within the same tumor. While the oncogenic role of the mutant allele is well established, the co-expressed WT ß-catenin, a key component of mechanotransduction pathways, may also influence the tumor cells. Thus, this work highlights interplay between WT and mutant ß-catenins through their differential regulation of *RND3* expression.

We first demonstrated that *RND3* expression is low in all subgroups of human HCCs and above all in ß-catenin-mutated tumors. Using cultured cells, we further found that silencing of ß-catenin led to an increase in *RND3* expression demonstrating that ß-catenin represses it. Interestingly, this effect was observed in both mutated or non-mutated cell lines. To dissect the individual contributions of WT and mutated ß-catenin to *RND3* expression, we used our dual ß-catenin knocked-down HepG2 model, which allows the specific silencing of either or both ß-catenin forms (WT/mutated). Using this model, we have previously identified new targets of ß-catenin in liver tumor cells, such as the actin-bunding protein, Fascin-1 and the small extracellular vesicle regulator, Rab27A (Gest et al., 2023)(Dantzer et al., 2024b). Here, we found that *RND3* expression is repressed by both the WT and the mutated forms of ß-catenin in HepG2 cells.

The WT pool of ß-catenin in HepG2 cells is mainly involved in structural activity by binding to E-cadherin in order to maintain cell-cell junctions. Indeed, we previously showed that the removal of WT ß-catenin disrupts E-cadherin-based junctions (Gest et al., 2023). These junctions, which are connected to the actin cytoskeleton, function as mechanosensors able of initiating signaling cascades that influence many aspects of cell behavior including cell fate, migration or division (Mammoto and Ingber, 2009). This occurs mainly through the regulation of gene expression after modulation of transcription factor activity. Among the key mechanotransduction pathways, the YAP/TAZ signaling cascade is intrinsically linked to the Wnt/ß-catenin pathway. We indeed found that the knockdown of WT ß-catenin alters this mechano-transduction pathway, resulting in increased expression of TEAD transcriptional targets. Our results show that WT ß-catenin regulates *RND3* expression *via* its promoter through the involvement of YAP and TEAD. Supporting this, YAP overexpression strongly increased *RND3* mRNA levels demonstrating that *RND3* is a target of this mechanosensitive pathway. These data obtained in cell lines were corroborated in human HCC samples, where *RND3* expression positively correlated with TEAD target gene expression. This regulation may occur in various cancers as *RND3* has also been described as an Hippo pathway target in colorectal cancers in response to the histone demethylase KDM3A activation (Wang et al., 2019). Additionally, we previously found that *RND3* is regulated by the mechano-responsive MAL/SRF pathway in HCC (Piquet et al., 2018). Altogether, these data establish *RND3* as a mechano-activable gene responsive to multiple mechano-stimuli such as cell-cell junction modification and actin cytoskeleton reorganization.

Our data also demonstrated that *RND3* expression is repressed by the oncogenic form of ß-catenin, which is characterized by uncontrolled transcriptional activity. In an unexpected manner, we demonstrated that this regulation occurs through *RND3* 3’UTR, revealing a novel post-transcriptional function of ß-catenin beyond its canonical transcriptional roles described (Sevim et al., 2025). The challenge is to distinguish between β-catenin’s well-established transcriptional role and this additional post-transcriptional regulation. Using our dual ß-catenin knocked-down HepG2 model, we obtained evidence supporting post-transcriptional regulation of *RND3* by ß-catenin. By cross-analyzing, miRNAs downregulated upon mutated ß-catenin silencing with those miRNAs predicted to bind the *RND3* 3’UTR, we indeed identified the miR-512-3p as a key regulator. Consistent with this post-transcriptional regulation, *RND3* and miR-512-3p negatively correlates in human HCC samples.

Thus, we have identified two new ways of regulation of *RND3*, an atypical member of the Rho GTPase family: at its transcriptional level *via* YAP/TEAD pathway and at its post-transcriptional level through miR-512-3p, both under the control of the dual function of ß-catenin. This study highlights the complexity of regulation that occurs upon ß-catenin expression and activation in liver cells. As a novel target of ß-catenin, Rnd3 may be involved in both its physiological and pathological roles in the liver. Rnd3 may constitute a key protein involved in the transcriptional program driven by oncogenic ß-catenin in HCC as well as in the mechanosensitive response linked to cell-cell adhesive communication in hepatocytes.

## List of Abbreviations

GS: Glutamine Synthetase
HCC: hepatocellular carcinoma

## ACKNOWLEDGEMENTS

We thank the OneCell and Vect’UB facilities (TBMCore, UMS005, Bordeaux) for the help with qRT-PCR and lentivirus experiments. Small RNA sequencing was performed at the PGTB (Univ. Bordeaux, INRAE, BIOGECO, F-33610 Cestas, France) with the help of Zoé Delporte. We warmly thank Pr. Jessica Zucman-Rossi and Gabrielle Couch (Cordeliers Research Center, Inserm, Paris Cite University, France) for their help in analyzing *RND3* expression n in human HCC.

