## Supplementary figures and images for "Wild-type and mutated ß-catenin differently repress *RND3/RHOE* expression in hepatocellular carcinoma"

### Supplemental figures

#### Supplemental Figure 1

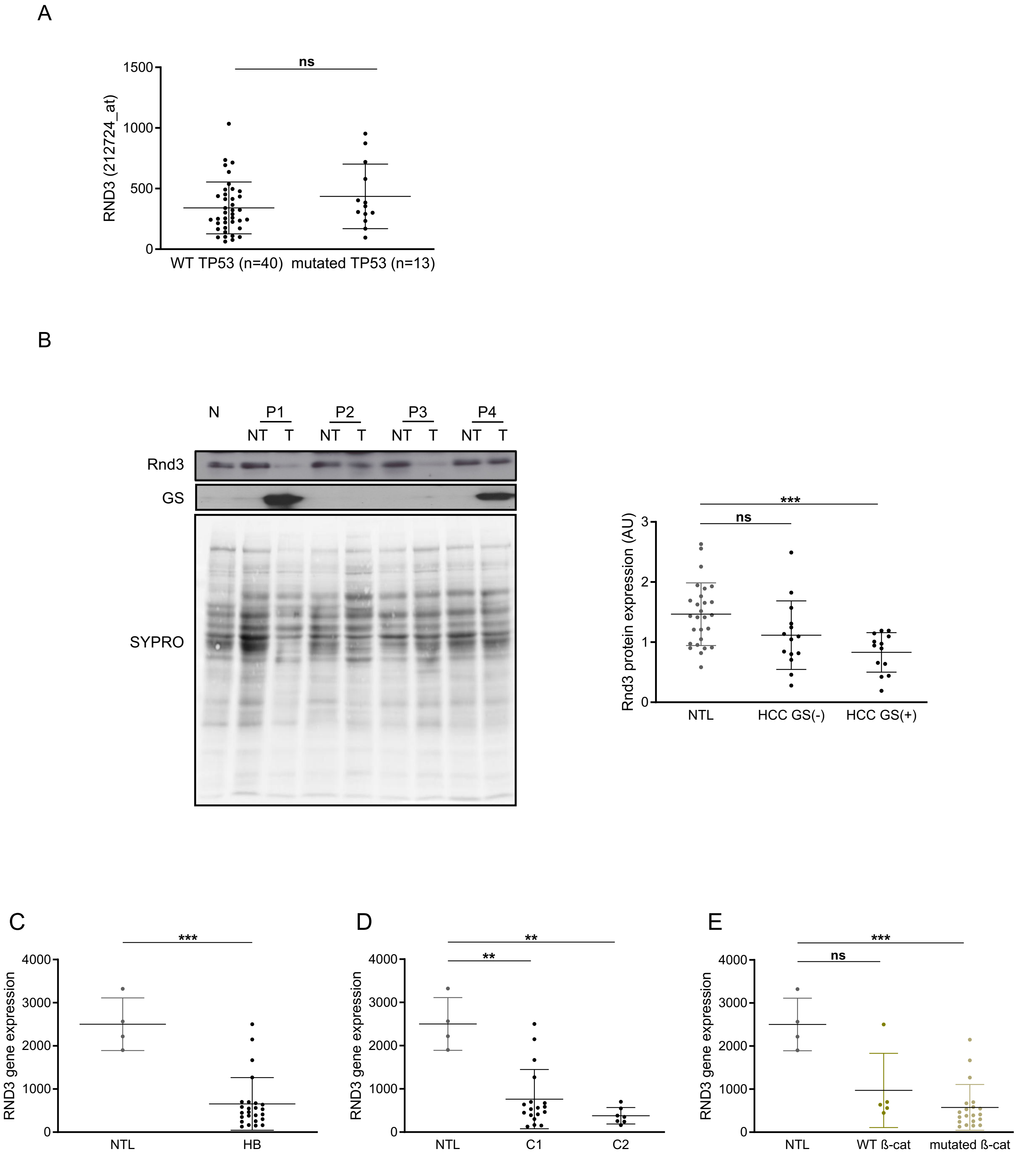

#### Supplemental Figure 2

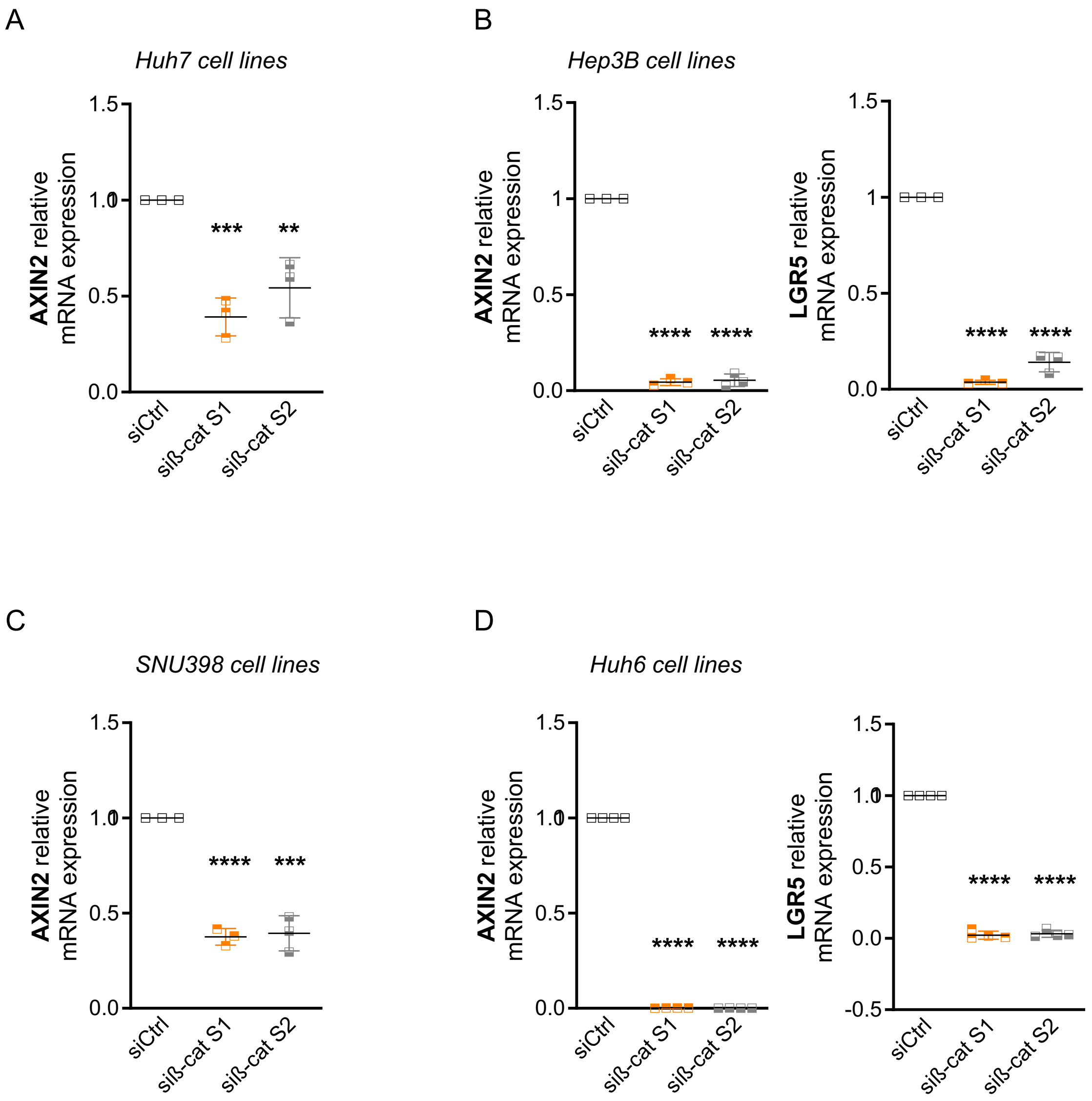

#### Supplemental Figure 3

A

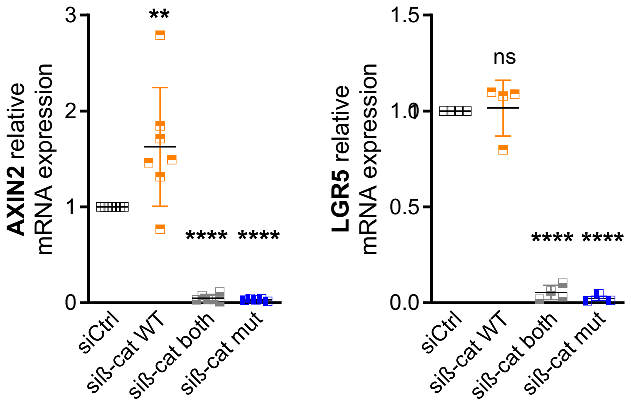

#### Supplemental Figure 4

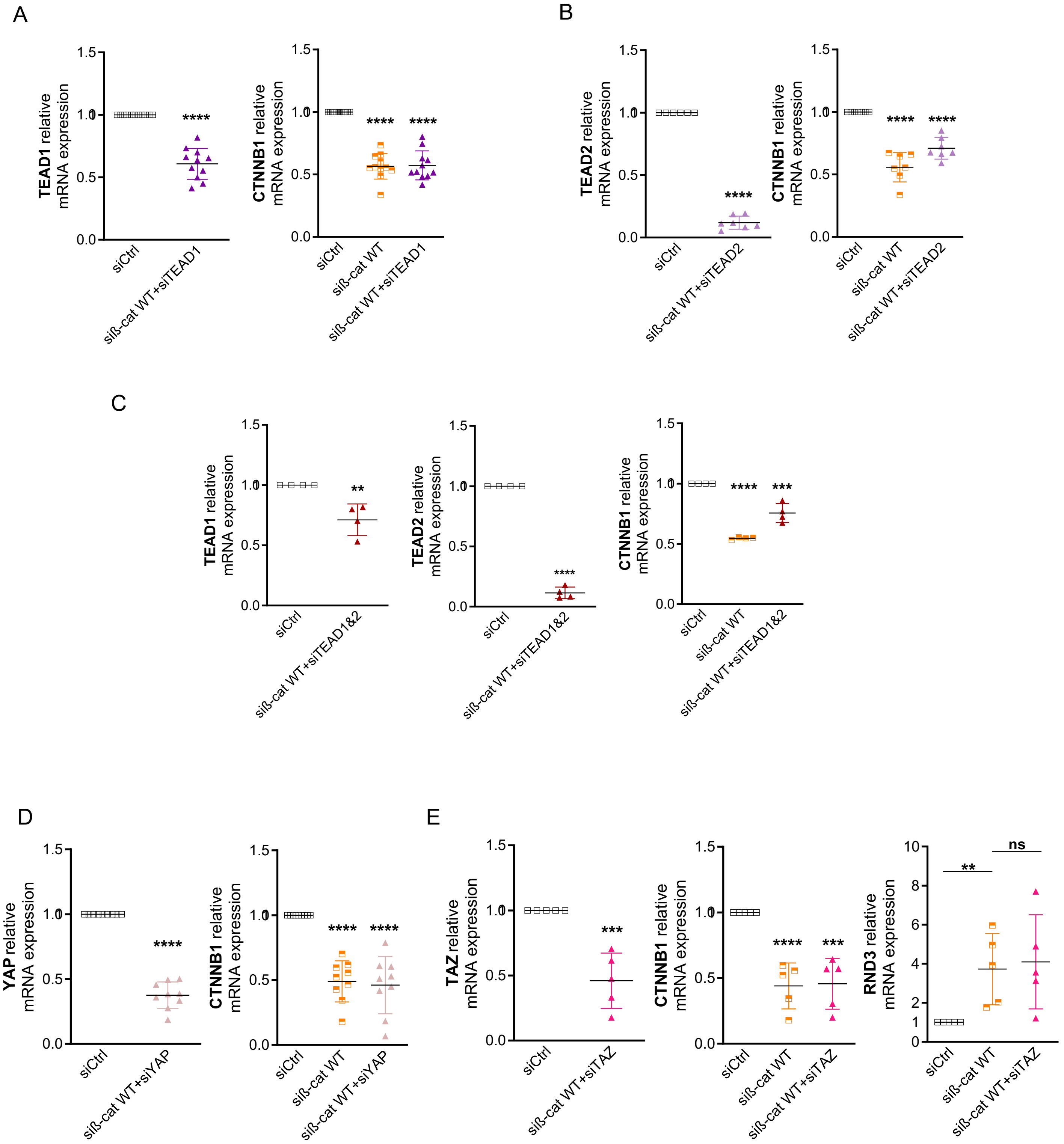
