## Supplemental material for "Wild-type and mutated ß-catenin differently repress *RND3/RHOE* expression in hepatocellular carcinoma"

**SUPPLEMENTARY MATERIAL**

**Supplemental figure 1:** ***RND3* expression is downregulated in human HCC and in hepatoblastoma**

**(A)** *RND3* gene expression in human HCC with wild-type versus mutated TP53 from the Boyault et al. cohort. **(B)** Rnd3 and glutamine synthase (GS) protein expression in HCC patient samples, showing tumor (T) and matched non-tumor (NT, n=27 patients total) tissues from the same patients. SYPRO staining was used to detect the total protein loaded. Patients were grouped based on GS expression in the tumor (GS(+); n=13 or GS(-); n=14). The graph on the right shows Rnd3 protein quantification in tumors with GS(+) and GS(-) patients, compared to Rnd3 levels in their matched non-tumor liver (NTL) tissues. The blot on the left is an example in 4 paired cases **(C)** Transcriptomic analysis from Cairo et al. showing *RND3* gene expression in non-tumor liver (NTL, n=4) tissue and hepatoblastoma (HB, n=25) tumors. **(D)** Same patients as in (C), classified into C1 (n=18) and C2 (n=7) molecular groups. **(E)** Same cohort classified according to ß-catenin mutation status (WT, n=5; mutated, n=20). Results are expressed as Mean ±SD, Student's t-test. ns for not significant; ***P*<0.01; ****P*<0.001.

**Supplemental figure 2:** **Validation of the ß-catenin inhibition in HCC cell lines**

Expression of ß-catenin target genes *AXIN2* and/or *LGR5* was assessed by qRT-PCR after siRNA-mediating silencing of ß-catenin (S1 and S2). Validation was performed in the following cell lines: Huh7 (A), Hep3B (B), SNU398 (C) and Huh6 (D). Results are representative of at least three independent experiments and expressed as Mean ±SD, Student's t-test. ***P*<0.01; ****P*<0.001; *****P*<0.0001.

**Supplemental figure 3:** **Confirmation of mutated ß-catenin inhibition in HepG2 cells.**

**(A)** Expression levels of the ß-catenin target genes *AXIN2* and *LGR5* were assessed by qRT-PCR after transfection of HepG2 cells with siRNAs targeting the WT (siß-cat WT), mutated (siß-cat mut) or both (siß-cat both) forms of ß-catenin. Results are representative of at least three independent experiments and expressed as Mean ±SD, Student's t-test. ns for not significant; **P*<0.05; *****P*<0.0001.

**Supplemental figure 4: Evaluation of WT ß-catenin knockdown and YAP/TAZ/TEAD expression in HepG2 cells.**

**(A-D)** After transfection of HepG2 cells with siRNA targeting the WT form of ß-catenin with or without siRNA against TEAD1 (A) or TEAD2 (B) or both (C), or YAP (D), the inhibition was confirmed by analyzing the expression of WT ß-catenin (*CTNNB1*) and *TEAD1* (A) or *TEAD2* (B) or both (C) or *YAP* (D). **(E)** HepG2 cells were transfected with siRNA against WT form of ß-catenin in the presence or absence of siRNA targeting TAZ and the expression of *TAZ*, *CTNNB1* and *RND3* were evaluated. Results are representative of at least three independent experiments and expressed as Mean ±SD, Student's t-test. ns for not significant; **$P$<0.01; ***$P$<0.001; ****$P$<0.0001.
