## Supplemental Table 3 for "Wild-type and mutated ß-catenin differently repress *RND3/RHOE* expression in hepatocellular carcinoma"

| Name | Fold change | FDR p-value | P-value |
|---|---|---|---|
| hsa-miR-155-5p | 80,4556471 | 0,07197454 | 0,008764 |
| hsa-miR-375-3p | 24,054906 | 1,49E-22 | 1,93E-25 |
| hsa-miR-92a-2-5p | 17,7087019 | 0,003943 | 0,0001788 |
| hsa-miR-885-5p | 15,0273518 | 2,416E-13 | 6,258E-16 |
| hsa-miR-616-5p | 11,3205813 | 0,06385913 | 0,007315 |
| hsa-miR-885-3p | 10,1131877 | 9,66E-13 | 3,754E-15 |
| hsa-miR-1973 | 9,56127562 | 0,004453 | 0,0002076 |
| hsa-miR-548ar-5p | 9,24541018 | | 0,01330948 |
| hsa-miR-9902 | 7,17311497 | | 0,03045113 |
| hsa-miR-4485-3p | 6,73513614 | 0,001564 | 0,00006482 |
| hsa-miR-4745-3p | 6,00230162 | 0,06149498 | 0,006452 |
| hsa-miR-642a-5p | 5,76849879 | 0,10371864 | 0,01504726 |
| hsa-miR-449a | 5,40663834 | 1,107E-06 | 1,721E-08 |
| hsa-miR-504-3p | 5,34832875 | 0,06862633 | 0,008224 |
| hsa-miR-642a-3p | 5,20479098 | 0,00498 | 0,0002645 |
| hsa-miR-6777-5p | 4,84577814 | 0,0526524 | 0,004842 |
| hsa-miR-3614-5p | 4,54861811 | 5,153E-06 | 1,012E-07 |
| hsa-miR-204-3p | 4,54458676 | | 0,03437225 |
| hsa-miR-6501-5p | 4,41505127 | 0,10034959 | 0,0139839 |
| hsa-miR-363-5p | 4,093757 | 0,03559028 | 0,003042 |
| hsa-miR-187-3p | 3,9902746 | 0,00002307 | 6,275E-07 |
| hsa-miR-127-3p | 3,91781741 | 0,008853 | 0,000539 |
| hsa-miR-504-5p | 3,84894039 | 0,14938263 | 0,0266008 |
| hsa-miR-4518 | 3,77689457 | 0,00498 | 0,0002395 |
| hsa-miR-3614-3p | 3,72947002 | 0,0001028 | 2,929E-06 |
| hsa-miR-4775 | 3,59213443 | 0,11847561 | 0,01902976 |
| hsa-miR-204-5p | 3,5146002 | 0,06385913 | 0,00724 |
| hsa-miR-431-5p | 3,48291911 | 0,07274571 | 0,009046 |
| hsa-miR-493-3p | 3,4807923 | 0,15788195 | 0,02905725 |
| hsa-miR-432-5p | 3,47373469 | 0,12500892 | 0,02024108 |
| hsa-miR-449c-5p | 3,33024046 | 0,22377686 | 0,04956716 |
| hsa-miR-664b-3p | 3,20208965 | 0,009399 | 0,0005844 |
| hsa-miR-22-5p | 3,17279405 | 9,091E-07 | 1,295E-08 |
| hsa-miR-215-5p | 2,97614232 | 0,06769298 | 0,007979 |
| hsa-miR-152-3p | 2,95025985 | 0,01732223 | 0,001144 |
| hsa-miR-196a-5p | 2,90939714 | 0,00498 | 0,0002581 |
| hsa-miR-363-3p | 2,78213059 | 0,02172593 | 0,001541 |
| hsa-miR-20b-5p | 2,7777473 | 0,007069 | 0,0003938 |
| hsa-miR-5591-5p | 2,75573433 | 0,16448522 | 0,03096469 |
| hsa-miR-125b-5p | 2,61656346 | 0,06142359 | 0,006176 |
| hsa-miR-153-3p | 2,54332095 | 0,03559028 | 0,003018 |
| hsa-miR-483-3p | 2,52396584 | 0,0001319 | 3,929E-06 |
| hsa-miR-664a-3p | 2,49597652 | 0,05535768 | 0,005235 |
| hsa-miR-99a-5p | 2,42379939 | 0,06326066 | 0,006965 |
| hsa-miR-6501-3p | 2,3481999 | 0,10237081 | 0,01458651 |

| | | | |
|---|---|---|---|
| hsa-miR-22-3p | 2,34041126 | 0,00001721 | 4,013E-07 |
| hsa-miR-3613-5p | 2,29657218 | 0,10719431 | 0,01596806 |
| hsa-miR-126-3p | 2,29463312 | 0,00498 | 0,000259 |
| hsa-miR-378i | 2,23157775 | 0,0065 | 0,0003536 |
| hsa-miR-378d | 2,18798487 | 0,03543262 | 0,002846 |
| hsa-miR-320e | 2,14972931 | 0,22136923 | 0,04731337 |
| hsa-miR-122-5p | 2,12069725 | 0,002774 | 0,0001222 |
| hsa-miR-143-3p | 2,10633411 | 0,00002307 | 6,159E-07 |
| hsa-let-7c-5p | 2,0768124 | 0,11343069 | 0,01763171 |
| hsa-miR-628-5p | 2,01697855 | 0,06326066 | 0,006921 |
| hsa-miR-1250-5p | 1,98375878 | 0,14793086 | 0,02595964 |
| hsa-miR-5699-5p | 1,95823461 | 0,22377686 | 0,04937776 |
| hsa-miR-192-5p | 1,94306508 | 0,0004501 | 0,00001691 |
| hsa-miR-628-3p | 1,9271189 | 0,19723845 | 0,04113392 |
| hsa-miR-10a-5p | 1,90770505 | 0,22223163 | 0,04864915 |
| hsa-miR-194-5p | 1,90480841 | 0,000412 | 0,00001494 |
| hsa-miR-664a-5p | 1,89583314 | 0,11314954 | 0,01722492 |
| hsa-miR-499a-5p | 1,87946365 | 0,14793086 | 0,02606036 |
| hsa-miR-483-5p | 1,86753421 | 0,03350993 | 0,002604 |
| hsa-miR-145-3p | 1,85237543 | 0,08842605 | 0,01179778 |
| hsa-miR-338-3p | 1,85047538 | 0,00498 | 0,0002583 |
| hsa-miR-126-5p | 1,82199137 | 0,07721152 | 0,01010151 |
| hsa-miR-378a-5p | 1,82192985 | 0,14938263 | 0,02670311 |
| hsa-miR-1301-3p | 1,81155409 | 0,08025857 | 0,01060411 |
| hsa-miR-194-3p | 1,76784357 | 0,19649184 | 0,04021465 |
| hsa-miR-548w | 1,7511261 | 0,22223163 | 0,04832287 |
| hsa-miR-378a-3p | 1,74997737 | 0,008395 | 0,0005002 |
| hsa-miR-12136 | 1,72406741 | 0,18577809 | 0,03657807 |
| hsa-miR-224-5p | 1,59659745 | 0,03559028 | 0,002948 |
| hsa-miR-130a-3p | 1,58213758 | 0,19649184 | 0,04012855 |
| hsa-miR-345-5p | 1,56052008 | 0,06142359 | 0,006206 |
| hsa-miR-574-5p | 1,5337359 | 0,18577809 | 0,0364096 |
| hsa-miR-1269b | 1,48499781 | 0,0615989 | 0,006552 |
| hsa-miR-146b-5p | 1,45800709 | 0,18637633 | 0,03693728 |
| hsa-miR-26b-5p | 1,45443181 | 0,10034959 | 0,01416853 |
| hsa-miR-10b-5p | 1,45233943 | 0,18537544 | 0,03601854 |
| hsa-miR-140-3p | 1,39368507 | 0,19600572 | 0,03960737 |
| hsa-miR-29a-3p | 1,38408928 | 0,13864821 | 0,0231679 |
| hsa-miR-320c | 1,37716671 | 0,21554101 | 0,04550931 |
| hsa-miR-671-5p | 1,35235885 | 0,22084896 | 0,0469161 |

| Name | Fold change | FDR p-value | P-value |
|---|---|---|---|
| hsa-miR-1283 | -20,586122 | 1,037E-09 | 5,373E-12 |
| hsa-miR-516a-5p | -18,149654 | 6,035E-08 | 3,909E-10 |
| hsa-miR-486-3p | -17,396834 | 0,00000253 | 4,588E-08 |
| hsa-miR-520b-5p | -16,295949 | 9,091E-07 | 1,293E-08 |
| hsa-miR-519a-3p | -13,211976 | 7,797E-08 | 6,06E-10 |
| hsa-miR-522-3p | -13,075675 | 1,537E-07 | 1,394E-09 |
| hsa-miR-519c-5p | -11,041932 | 1,652E-07 | 1,712E-09 |
| hsa-miR-486-5p | -9,0352605 | 2,713E-07 | 3,163E-09 |
| hsa-miR-23a-5p | -7,3063806 | 0,02001017 | 0,001374 |
| hsa-miR-549a-5p | -6,4814671 | 0,000209 | 7,039E-06 |
| hsa-miR-135a-3p | -4,5929453 | 0,0908609 | 0,01224033 |
| hsa-miR-519c-3p | -3,8664916 | | 0,03803415 |
| hsa-miR-3154 | -3,8052206 | 0,11314954 | 0,01729488 |
| hsa-miR-1180-5p | -3,7091689 | 0,13550137 | 0,02229103 |
| hsa-miR-515-3p | -3,6442621 | 0,01790007 | 0,001206 |
| hsa-miR-181c-3p | -3,6409952 | 0,0009233 | 0,00003707 |
| hsa-miR-135a-5p | -3,6153846 | 0,06688763 | 0,007798 |
| hsa-miR-181d-5p | -3,6087413 | 0,000519 | 0,00002017 |
| hsa-miR-7109-5p | -3,5278748 | 0,18831304 | 0,03756504 |
| hsa-miR-524-5p | -3,4249291 | 0,10641864 | 0,01571467 |
| hsa-miR-222-5p | -3,3345294 | 0,00001994 | 4,908E-07 |
| hsa-miR-526b-5p | -3,3260628 | 0,06149498 | 0,006447 |
| hsa-miR-1909-3p | -3,2166374 | 0,19723845 | 0,04069578 |
| hsa-miR-219b-3p | -3,1980497 | 0,06862633 | 0,008267 |
| hsa-miR-615-3p | -3,1661302 | 0,11813686 | 0,01882232 |
| hsa-miR-199b-5p | -3,1394875 | 0,000375 | 0,00001311 |
| hsa-miR-512-3p | -3,1295436 | 0,05617741 | 0,005385 |
| hsa-miR-3929 | -3,1058914 | 0,03164344 | 0,002418 |
| hsa-miR-518f-3p | -3,067509 | 0,07313724 | 0,009284 |
| hsa-miR-4726-5p | -3,05538 | 0,05985949 | 0,005893 |
| hsa-miR-520a-5p | -3,0410261 | 0,02336294 | 0,001695 |
| hsa-miR-199a-3p | -2,9393771 | 0,0001837 | 0,00000571 |
| hsa-miR-365a-5p | -2,9101922 | 0,007172 | 0,0004161 |
| hsa-miR-520f-3p | -2,8794006 | 0,03768784 | 0,003368 |
| hsa-miR-6515-5p | -2,8227418 | 0,06385913 | 0,007231 |
| hsa-miR-3170 | -2,7503596 | 0,18491206 | 0,03568899 |
| hsa-miR-135b-3p | -2,743915 | 1,327E-06 | 2,234E-08 |
| hsa-miR-498-5p | -2,7403776 | 0,10034959 | 0,01406295 |
| hsa-miR-142-5p | -2,7293909 | 0,03760406 | 0,003312 |
| hsa-miR-454-5p | -2,6662289 | 0,12886201 | 0,02103188 |
| hsa-miR-519d-3p | -2,6411671 | 0,14068361 | 0,02387248 |
| hsa-miR-373-5p | -2,6270042 | 0,19723845 | 0,04093889 |
| hsa-miR-181c-5p | -2,5928629 | 0,02172593 | 0,001548 |
| hsa-miR-193b-3p | -2,516286 | 0,05274317 | 0,004919 |
| hsa-miR-4523 | -2,4905968 | 0,05741632 | 0,005578 |

| | | | |
|---|---|---|---|
| hsa-miR-549a-3p | -2,4754586 | 0,02963844 | 0,002188 |
| hsa-miR-27a-5p | -2,4751589 | 0,06148708 | 0,006292 |
| hsa-miR-142-3p | -2,4677759 | 0,04838665 | 0,004387 |
| hsa-let-7e-3p | -2,4128161 | 0,06385913 | 0,007362 |
| hsa-miR-372-5p | -2,3805918 | 0,14063528 | 0,02368211 |
| hsa-miR-146a-5p | -2,355882 | 0,0001935 | 6,265E-06 |
| hsa-miR-515-5p | -2,2895893 | 0,11813686 | 0,01877023 |
| hsa-miR-6783-3p | -2,2847035 | 0,07313724 | 0,009227 |
| hsa-miR-193b-5p | -2,2781254 | 0,0615989 | 0,006623 |
| hsa-miR-1323 | -2,2720178 | 0,1164607 | 0,01825356 |
| hsa-miR-4796-3p | -2,2542026 | 0,09959445 | 0,01367489 |
| hsa-miR-27a-3p | -2,2472264 | 0,01055522 | 0,00067 |
| hsa-miR-144-3p | -2,2191771 | 0,1549828 | 0,02790493 |
| hsa-miR-221-3p | -2,1997557 | 5,153E-06 | 1,068E-07 |
| hsa-miR-518c-3p | -2,1141951 | 0,18870097 | 0,03788685 |
| hsa-miR-5580-3p | -2,1061608 | 0,09944543 | 0,01352561 |
| hsa-miR-4747-5p | -2,0950552 | 0,10371864 | 0,01501897 |
| hsa-miR-942-3p | -2,0918256 | 0,16246192 | 0,03030378 |
| hsa-miR-455-5p | -2,0655678 | 0,03164344 | 0,0024 |
| hsa-miR-455-3p | -2,0626326 | 0,00001721 | 3,794E-07 |
| hsa-miR-4699-5p | -2,0588392 | 0,15788195 | 0,02924497 |
| hsa-miR-23a-3p | -2,0245066 | 0,03708139 | 0,003218 |
| hsa-miR-373-3p | -2,0182036 | 0,10551727 | 0,01544489 |
| hsa-miR-372-3p | -1,9796285 | 0,14348407 | 0,02490527 |
| hsa-miR-7974 | -1,8975281 | 0,20663079 | 0,04336035 |
| hsa-miR-365a-3p | -1,8960306 | 0,16938804 | 0,03225394 |
| hsa-miR-550a-3p | -1,8251048 | 0,22223163 | 0,04858001 |
| hsa-miR-15b-5p | -1,8200012 | 0,002641 | 0,0001129 |
| hsa-miR-135b-5p | -1,7960898 | 0,03559028 | 0,003043 |
| hsa-miR-629-5p | -1,7541813 | 0,11026609 | 0,01656848 |
| hsa-miR-1296-5p | -1,7465598 | 0,15656043 | 0,02859459 |
| hsa-miR-429 | -1,6960112 | 0,007172 | 0,0004181 |
| hsa-miR-24-3p | -1,6109834 | 0,01073438 | 0,0006952 |
| hsa-miR-200a-3p | -1,5552657 | 0,07553824 | 0,009687 |
| hsa-miR-4455 | -1,5522664 | 0,03466189 | 0,002739 |
| hsa-miR-15a-5p | -1,513881 | 0,15656043 | 0,0285833 |
| hsa-miR-93-3p | -1,5050938 | 0,07210063 | 0,008872 |
| hsa-miR-16-5p | -1,4757226 | 0,07585343 | 0,009826 |
| hsa-miR-744-5p | -1,4615779 | 0,14348407 | 0,02478819 |
| hsa-miR-20a-3p | -1,4350087 | 0,16448522 | 0,03110731 |
| hsa-miR-1304-3p | -1,4207159 | 0,22223163 | 0,04864806 |
| hsa-miR-130b-3p | -1,4097924 | 0,11343069 | 0,01755018 |
| hsa-miR-107 | -1,4041319 | 0,14200099 | 0,02427996 |
| hsa-miR-103a-3p | -1,3917811 | 0,13864821 | 0,02309364 |
| hsa-miR-106b-5p | -1,3518771 | 0,18394733 | 0,03526451 |
