## Supplemental Table 2 for "Wild-type and mutated ß-catenin differently repress *RND3/RHOE* expression in hepatocellular carcinoma"

### Supplemental table 2: Primer sequences for real-time quantitative PCR

| *Genes* | *Primer sequences* |
|---|---|
| ***AMOTL2*** | F: AGTGAGCGACAAACAGCAGACG<br>R: ATCTCTGCTCCCGTGTTTGGCA |
| ***ANKRD*** | F: AGACTCCTTCAGCCAACATGATG<br>R: CTCTCCATCTCTGAAATCCTCAGG |
| ***ARHGAP29*** | F: TTGCAGCTCTCCAGGCTAAC<br>R: AGATGCTCCTCTTCTGCACG |
| ***AXIN2*** | F: TGCTCTGTTTTGTCTTAAAGGTCTTGA<br>R: ACAGATCATCCCATCCAACACA |
| ***AXL*** | F: CGTAACCTCCACCTGGTCTC<br>R: TCCCATCGTCTGACAGCA |
| ***CTNNB1*** | F: TCTTACACCCACCATCCCAC<br>R: GCACGAACAAGCAACTGAAC |
| ***CYR61*** | F: ATGAATTGATTGCAGTTGGAAA<br>R: TAAAGGGTTGTATAGGATGCGA |
| ***LGR5*** | F: GAGGATCTGGTGAGCCTGAGAA<br>R: GATGCTGGAGCTGGTAAAGGT |
| ***MST1*** | F: CCTCCCACATTCCGAAAACCA<br>R: GCACTCCTGACAAATGGGTG |
| ***MST2*** | F: TTTCCTCCTTTCGGCCTTCG<br>R: GCGACAACTTGACCGGATTC |
| ***RND3*** | F: GCCAGCCAGAAATTATCCAGC<br>R: CTTGGCGAAGACATGGAGC |
| ***TEAD1*** | F: CTGAGTCGCAGTTACCACCA<br>R: AGCCTGGAGCCTTTTCAAG |
| ***TEAD2*** | F: ACATGATGAACAGCGTCCTG<br>R: CAGCAGTTCCTGGGTGTCTC |
| ***YAP*** | F: CACAGCATGTTCGAGCTCAT<br>R: GATGCTGAGCTGTGGGTGTA |
| ***TAZ*** | F: GTTTATGGGAGAGTCCGGGAG<br>R: AGTCTAAGGGCTTCGGCTCT |
