## Supplemental Table 1 for "Wild-type and mutated ß-catenin differently repress *RND3/RHOE* expression in hepatocellular carcinoma"

### Supplemental Table 1: siRNAs or miRNA sequences

| *siRNAs* | *Targeted sequences* | *References* |
|---|---|---|
| **siCtrl** | Not provided | Qiagen (AllStars Negative control) |
| **Siß-cat WT or S1** | GTAGCTGATATTGATGGACAG | Gest et al., 2023 |
| **Siß-cat Both or S2** | ACCAGTTGTGGTTAAGCTCTT | Gest et al., 2023 |
| **Siß-cat mut** | TGTTAGTCACAACTATCAAGA | Gest et al., 2023 |
| **SiTEAD1** | Not provided | Ambion #s13961 |
| **SiTEAD2** | Not provided | Ambion #s16075 |
| **siYAP** | GACATCTTCTGGTCAGAGA | Eurofins |
| **siTAZ** | ACGTTGACTTAGGAACTTT | Eurofins |
| **siAPC** | SMART pool | Dharmacon #L-009869 |
| **siCtrl** (for SMART pool) | Non-targeting pool | Dharmacon #D-001810 |

| *miRNAs* | *Mature sequence (guide strand)* | *References* |
|---|---|---|
| **hsa-miR-512-3p** | AAGUGCUGUCAUAGCUGAGGUC | Qiagen |
